# Heterogeneous Contribution of Microdeletions in the Development of Common Generalized and Focal epilepsies

**DOI:** 10.1101/131359

**Authors:** Eduardo Pérez-Palma, Ingo Helbig, Karl Martin Klein, Verneri Anttila, Heiko Horn, Eva Maria Reinthaler, Padhraig Gormley, Andrea Ganna, Andrea Byrnes, Katharina Pernhorst, Mohammad R. Toliat, EuroEPINOMICS-RE consortium, Italian League against Epilepsy Consortium, Elmo Saarentaus, Daniel P. Howrigan, Per Hoffman, Juan Francisco Miquel, Giancarlo De Ferrari, Peter Nürnberg, Holger Lerche, Fritz Zimprich, Bern A. Neubauer, Albert J. Becker, Felix Rosenow, Emilio Perucca, Federico Zara, Yvonne G. Weber, Dennis Lal

## Abstract

**Background:** Microdeletions are known to confer risk to epilepsy, particularly at genomic rearrangement “hotspot” loci. However, deciphering their role outside hotspots and risk assessment by epilepsy sub-type has not been conducted.

**Methods:** We assessed the burden, frequency and genomic content of rare, large microdeletions found in a previously published cohort of 1,366 patients with Genetic Generalized Epilepsy (GGE) plus two sets of additional unpublished genome-wide microdeletions found in 281 Rolandic Epilepsy (RE) and 807 Adult Focal Epilepsy (AFE) patients, totaling 2,454 cases. These microdeletion sets were assessed in a combined analysis and in sub-type specific approaches against 6,746 ethnically matched controls.

**Results:** When hotspots are considered, we detected an enrichment of microdeletions in the combined epilepsy analysis (adjusted-P= 2.00×10^-7^; OR = 1.89; 95%-CI: 1.51-2.35), where the implicated microdeletions overlapped with rarely deleted genes and those involved in neurodevelopmental processes. Sub-type specific analyses showed that hotspot deletions in the GGE subgroup contribute most of the signal (adjusted-P = 1.22×10^-12^; OR = 7.45; 95%-CI = 4.20-11.97). Outside hotspot loci, microdeletions were enriched in the GGE cohort for neurodevelopmental genes (adjusted-P = 4.78×10^-3^; OR = 2.30; 95%-CI = 1.42-3.70), whereas no additional signal was observed for RE and AFE. Still, gene content analysis was able to identify known (*NRXN1*, *RBFOX1* and *PCDH7*) and novel (*LOC102723362*) candidate genes affected in more than one epilepsy sub-type but not in controls.

**Conclusions:** Our results show a heterogeneous effect of recurrent and non-recurrent microdeletions as part of the genetic architecture of GGE and a minor to negligible contribution in the etiology of RE and AFE.

## INTRODUCTION

Epilepsies comprise a clinically complex group of neurological disorders defined by recurrent spontaneous seizures[1], with an age-adjusted global prevalence estimated in the range of 2.7-17.6 per 1,000 individuals[2]. The most common types of epilepsies represent the heterogeneous group of acquired and non-acquired Adult Focal Epilepsies (AFE), idiopathic Genetic Generalized Epilepsies (GGE), and idiopathic childhood focal Rolandic Epilepsies (RE). The genetic architecture of these common epilepsies is presumed to be complex as it has been described by a wide range of syndrome specific variant associations, as well as a few shared seizure susceptibility variants (for review see [3]). Twin studies have shown strong but differential concordance rates among epilepsies, including GGE and AFE with ∼80% and ∼40%, respectively[4 5], while the genetic contribution to RE may be related to the underlying electroencephalogram (EEG) pattern rather than the seizures themselves[6]. Despite these differences, familial enrichment for seizure disorders has been demonstrated and genetic risk factors in a single gene (*GRIN2A*) have recently been identified in > 7.5% of RE patients[7].

Copy number variants (CNVs) are genomic segments between 50 bp and 3 Mb in size which can behave as a loss and/or gain of genomic sequence relative to the reference genome based on the number of copies present[8]. CNVs are a significant source of genetic variation between two individuals and can, depending on their location on the chromosomes, cause changes in gene dosage, alternative splicing, or even lead to gene fusion events[9]. Microdeletions, defined as large (i.e. >400 kb) and rare CNVs (MAF < 1%) with a deletion behavior, are more likely to have damaging effects than duplications[8 10]. In addition, duplications are more prone to false positive calls as they are more difficult to identify in genotype array data.

Microdeletions are associated with a broad spectrum of neurological diseases such as autism (ASD)[11], schizophrenia (SCZ)[12] and intellectual disability (ID)[13]. In this regard, twelve genomic regions prone to exhibit copy number rearrangements or CNV “hotspot loci” have been reported to increase the risk for neurodevelopmental disorders[14], seven of them directly associated to epilepsy including 1q21.1, 15q11.2, 15q13.3, 16p11.2, 16p12, 16p13.11, and 22q11.2 to GGE[15–18], and also 16p11.2 to RE and AFE [19 20]. The relationship to epilepsy of the remaining 5 loci, namely 3q29, 10q22q23, 15q24, 17q12, and 17q21.3 remains to be elucidated. Overall, how these large and polygenic microdeletions increase risk for neurodevelopmental disorders is not fully understood. However, recent evidence from deletions in patients with GGE show an enrichment in genes involved in neurodevelopmental processes[18]. Whether this enrichment is also found in other types of common epilepsies has not yet been evaluated. Previous results support non-recurrent deletions in *RBFOX1*[20–22], *NRXN1*[23], and *GRIN2A[7 24]* in candidate gene studies in GGE/RE/AFE, GGE, and RE, respectively. However, a genome-wide comparison for shared or sub-type specific deleted genes in GGE, RE, and AFE has not been conducted.

Due to the low frequency of deletions, large sample sizes are required to identify novel susceptibility genes and to decipher syndrome-specific patterns. Considering that previous microdeletion associations are generally reported only within a particular type of epilepsy, our goal was to evaluate the global and specific contribution of microdeletions across GGE, RE, and AFE. Thus to investigate the deletion burden landscape for common types of epilepsy, here we present the largest combined microdeletion study to date for these traits, examining a total of 9,200 individuals (2,454 epilepsy cases and 6,746 controls). We combined the microdeletions found in a published GGE cohort with unpublished genome-wide microdeletions found in RE[19] and AFE[20] studies. We investigated the combined and sub-type specific burden of microdeletions, as well as their frequency, distribution and gene content. Finally, we assessed the protein-protein interactions and tissue specific expression pattern of genes that were only deleted in patients.

## MATERIALS AND METHODS

### Case-control cohorts

All patient and control cohorts included in the analysis are of European origin, have been genetically matched with their respective controls and have been described previously in detail [18–20] (Table 1). The epilepsy subtype classifications were based on the terminology proposed by the Commission on Classification and Terminology of the International League against Epilepsy (ILAE)[1]. For an extended description of phenotypes, sample recruitment, genotyping and CNV calling see the Supplementary Information file. Briefly, the epilepsy cohort was composed by 1,366 GGE cases genotyped with the Genome-Wide Human SNP Array 6.0 platform (Affymetrix, Santa Clara, CA, USA)[18] plus 281 and 807 RE and AFE cases, respectively which were genotyped using the Human OmniExpress BeadChip platform (Illumina Inc., San Diego, CA, U.S.A.). The control set was composed of 5,234 samples extracted from the original GGE study and genotyped with the Genome-Wide Human SNP Array 6.0 platform (n= 5,234[18]) plus 1,512 controls extracted from the original RE study [19] and genotyped using the Human OmniExpress BeadChip platform, totaling 6,746 control individuals (Table 1).

**Table 1.** Datasets main features.

| **Epilepsy sub-type** | **Code** | **Cases** | **Controls** | **Array** |
| --- | --- | --- | --- | --- |
| Genetic Generalized Epilepsies | GGE | 1,366 | 5,234 | Affymetrix GenomeWide 6.0 |
| Rolandic Epilepsy | RE | 281 | 1,512 | Illumina Omni Express Array |
| Adult Focal Epilepsy | AFE | 807 | - | Illumina Omni Express Array |
| **Total** |  | 2,454 | 6,746 | 9,200 |

### Microdeletion calling

The software PennCNV was used as the CNV calling algorithm following author recommendations for the corresponding Affymetrix and Illumina arrays[25]. Microdeletions were defined as autosomal CNVs with deletion behavior spanning at least 400kb in both Affymetrix and Illumina arrays to enrich for likely pathogenic variants. Additionally, to ensure highly reliable calls we only considered microdeletions involving at least 200 probes for the Affymetrix 6.0 platform [18 26]. Considering that amount of probes of the Illumina Omni express arrays (n= ∼740,000 markers) was less than half (40%) of the Affymetrix 6.0 platform (∼1,850,000 markers), but at the same time less prone to false positives[26], a threshold of 100 markers was applied to the Illumina calls. Only rare microdeletions, defined by a cohort specific frequency below 1% were considered for further analysis. Microdeletion frequency was calculated considering only the control cohort (n= 6,746).

### Burden statistics

For the combined epilepsy dataset, we evaluated patient and control autosomal microdeletion burden using a binomial logistic regression model implemented in the R statistical software[27]. To account for possible bias introduced due to the different genotyping platforms, we adjusted for this factor in the regression by including “platform” as a covariate in the model. Corresponding odds ratios (ORs) and 95%-confidence intervals (C.I.) were estimated from the log-likelihood function, while the association p-values for the combined regression coefficients were calculated using a Wald’s test with 1 degree of freedom. For the epilepsy subtype analysis each group (GGE, RE, and AFE) was tested individually for deletion enrichment in comparison to platform-matched controls. GGE samples were compared to the 5,234 Affymetrix controls while RE and AFE samples were compared individually to the 1,512 Illumina controls (Table 1). The p-values, corresponding odds ratios (ORs) and 95%-confidence intervals were calculated with a two-sided Fisher’s exact test. For both the combined and the sub-type analyses, multiple testing correction was applied using the Bonferroni method considering 8 comparisons (see microdeletion subsets interrogated below) to obtain adjusted p-values. Adjusted two-sided p-values < 0.05 were considered significant.

### Microdeletions subsets interrogated

We evaluated burden enrichment within 8 microdeletions subsets: 1) All microdeletions; 2) microdeletions overlapping hotspot loci; 3) microdeletions outside hotspot loci; 4) microdeletions overlapping “constrained” genes; 5) microdeletions overlapping “neurodevelopmental” genes; 6) microdeletions overlapping “ASD-related” genes (Autism Spectrum Disorders); 7) microdeletions overlapping “DDG2P” genes; and 8) microdeletions overlapping “loss of function intolerant” genes. While known CNV hotspot loci were extracted from a previous microdeletion revision [14] (No. genes = 330). Here, for subset number 4) we define “constrained” genes as those not overlapped by a CNV of the CNV control map reported by Zarrei et al, 2015[8] which constitutes a curated version of the database of genomic variants (DGV) on healthy individuals (No. genes = 20.208). For subset 5), “neurodevelopmental” genes were extracted from the original GGE study[18] and defined based on literature and database queries[18 28] (No. genes = 1.559). For subset 6), “ASD-related” genes were extracted from Uddin et al, 2014[29] and defined as those enriched for deleterious exonic de novo mutations in comparison with healthy siblings (No. genes = 1.683); For subset 7) “DDG2P” genes were extracted from a curated list of genes reported to be associated with developmental disorders, compiled by clinicians as part of the DDD study[30] to facilitate clinical feedback of likely causal variants (No. genes = 294; https://decipher.sanger.ac.uk/ddd#ddgenes). For subset 8), “loss of function intolerant” genes were extracted from Lek et al, 2016 [31] and defined by having less than expected loss of function variants within the 60,706 unrelated individuals from the exome aggregation consortium (No. genes = 2.506). The complete gene-sets are available as a publicly available resource in https://github.com/dlal-group/gene-sets).

### Tissue specific expression analysis

Gene expression analysis was performed using GTEx project resource (http://www.gtexportal.org/home/; version3), which includes gene expression data of 42 tissues from 1,561 human samples. Filtered candidate genes were used as a query to evaluate significant enrichment of tissue specific expression against a background distribution derived from multiple permutations. Extended description of the implemented methodology is provided in the Supplementary Information File.

### Network analysis

The data was partitioned into five microdeletion input sets; a combined burden analysis (GGE+RE+AFE), GGE only, RE only, AFE only, and finally an additional control set consisting of all microdeletions from the controls. The Disease Association Protein-Protein Link Evaluator software (DAPPLE, available at https://genepattern.broadinstitute.org/) was run with the CNV regions as input, using 10,000 iterations for each set. Network enrichment was calculated with the hypergeometric test over MSigDB pathways[32] adjusting for genes actually present in the InWeb network.

## RESULTS

The analysis was performed in two stages: First, the combined autosomal burden analysis for the entire epilepsy cohort (GGE+RE+AFE) followed by epilepsy subtype burden analyses (GGE, RE and AFE independently). In both strategies we tested burden enrichment among eight microdeletions subsets: 1) all microdeletions; 2) microdeletions overlapping hotspot loci and 3) microdeletions outside hotspot loci. Hotspot loci included collectively 12 loci: 1q21.1, 3q29, 10q22q23, 15q11.2, 15q13.3, 15q24, 16p11.2, 16p12, 16p13.11, 17q12, 17q21.3, and 22q11.2[14]. Subsequently, considering epilepsy etiology and comorbidities, microdeletions were filtered to include: 4) microdeletions overlapping “constrained” genes; 5) microdeletions overlapping “neurodevelopmental” genes; 6) microdeletions overlapping “ASD-related” genes; 7) microdeletions overlapping “Developmental Disorders” genes (DDG2P); and finally 8) microdeletions overlapping “loss of function intolerant” genes[31].

### Microdeletion frequency: Combined burden analysis

To investigate the overall contribution of microdeletions in the etiology of common types of epilepsy, we combined all the published microdeletions found in the GGE cohort [18] with the genome-wide microdeletions identified in RE[19] and AFE[20] cohorts (Table 1). In total, we analyzed 134 microdeletions in 2,454 patients with epilepsy compared to 219 microdeletions in 6,746 controls. Only 24 microdeletions (6.74 %) were found in more than one sample, with a maximum frequency of 0.059 % (n = 4 out of 6,746 control samples) which was reached by only two microdeletions. Thus, all included microdeletions were rare (<1%) and mostly singletons (93.04 %). Genomic context and sample annotation of all microdeletions found in the combined epilepsy sample is provided in Supplementary Table S1.

We compared the frequency of cases with microdeletions against the frequency of controls with microdeletions, interrogating their burden while controlling for batch (i.e. platform) effects with a linear regression model (see Methods). Overall, 5.42 % of cases (n = 133) carried at least one microdeletion compared to 3.46 % (n = 234) in controls (adjusted P= 2.00 × 10^-7^; OR = 1.89; 95%-CI: 1.51-2.35). The number of individuals carrying at least two microdeletions did not differ significantly between cases (No. cases = 4 out of 2,454) and controls (No. controls= 7 out 6,746; P = 0.498; OR = 1.57; 95%-CI: 0.33-6.18), suggesting that enrichment in patients for single microdeletions was not due to an excessive number of microdeletions per patient, which may arise due to batch effects in CNV calling. Focusing on coding regions, the burden of microdeletions overlapping at least one RefSeq gene was similar to the overall signal (adjusted P = 4.02 × 10^-7^; OR = 1.90; 95%-CI: 1.51-2.40).

For the combined results, the most significant enrichment was observed for microdeletions overlapping one of the 12 hotspot loci (adjusted P = 1.99 × 10^-12^; OR = 6.99; 95%-CI: 4.2-11.97), which is in agreement with previous observations in the GGE cohort that considered only 7 hotspot loci[18]. In this regard, we decided to examine the contribution of microdeletion burden in epilepsy patients outside these known hotspot loci. Thus, a total of 58 microdeletions were filtered out from the analysis (Supplementary Table S1). The contribution of microdeletions inside these regions is substantial, and the overall enrichment did not reach significance after hotspot loci had been removed (adjusted P = 0.17; OR = 1.34; 95%-CI: 1.03-1.73).

### Microdeletion distribution: Epilepsy sub-type burden analysis

We hypothesized that microdeletions outside of the hot spot loci are also conferring risk for the disease but are more heterogeneously distributed across the genome and epilepsy subtype specific. In this regard, we subsequently compared the microdeletion burden including hotspot loci for each epilepsy sub-type (GGE, AFE, and RE, Supplementary Figure S2) and then investigated whether we can identify enrichment for candidate microdeletion subsets not overlapping these regions (figure 1). As expected, the analysis showed that GGE patients were most significantly enriched for microdeletions overlapping hotspot loci (adjusted P = 1.22 × 10^-12^; OR = 7.45; 95%-CI: 4.21-13.5) and overlapping neurodevelopmental genes as previously shown in [18]. Interestingly, microdeletions overlapping neurodevelopmental genes not overlapping hotspot loci remained significant (adjusted P = 1.12 × 10^-2^; OR = 2.85; 95%-CI: 1.62-4.94). In contrast, RE patients showed nominal significance and a large effect size of microdeletions overlapping hotspot loci but this did not survive multiple testing correction (figure 1; nominal P = 0.029; adjusted P = 0.24; OR = 8.13; 95%-CI: 0.92-97.79). AFE patients did not show significant differences with control samples within hotspot loci (P = 1; OR = 2.85; 95%-CI: 0.13-25.9). Furthermore, RE and AFE samples were not enriched for any of the microdeletion subsets interrogated outside hotspot loci. Sample count of all the microdeletion subsets evaluated is provided in Supplementary Table S3.

**Figure 1.**
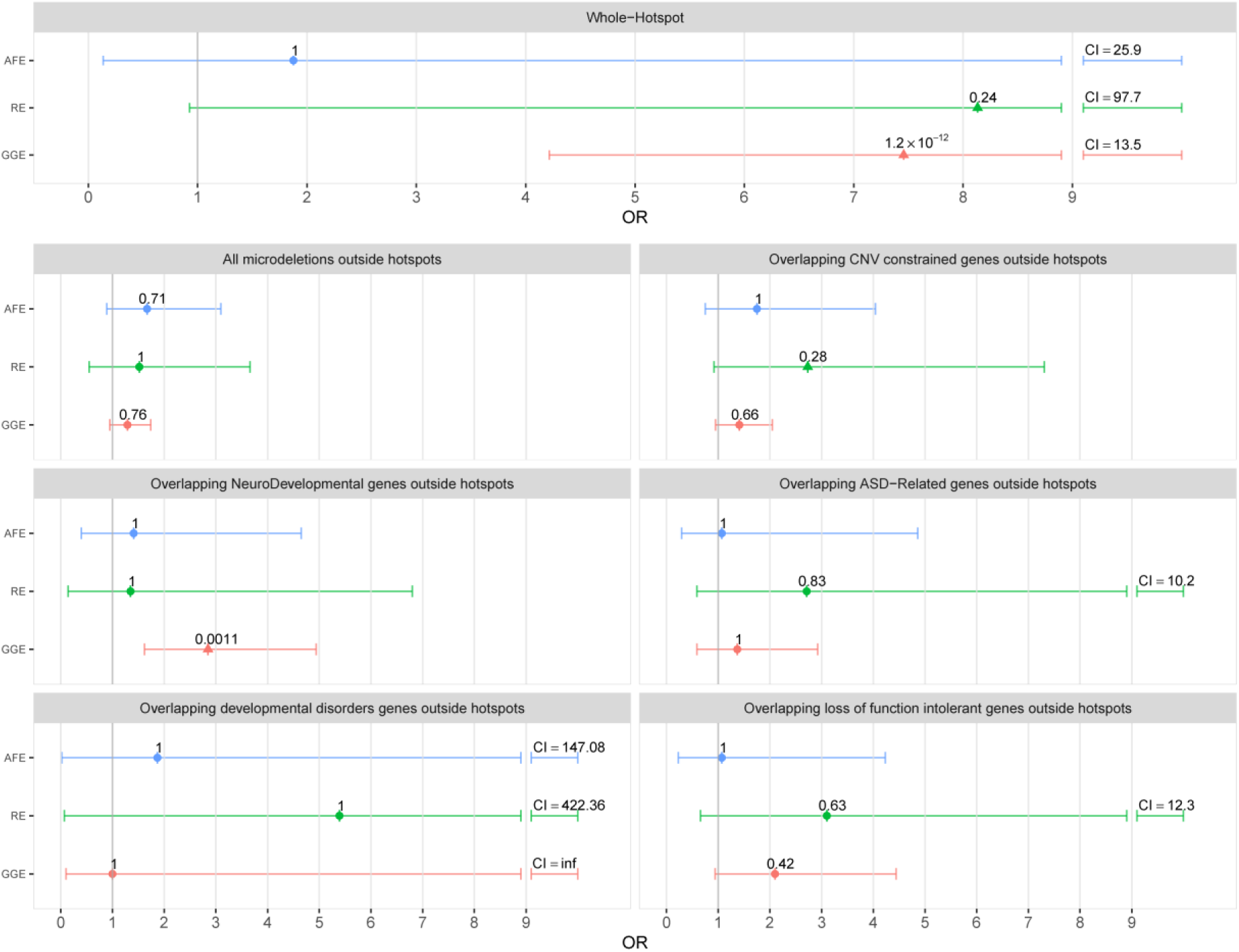
Microdeletion burden analysis in common epilepsy types. For adult focal epilepsies (AFE, cyan), rolandic epilepsies (RE, green), genetic generalized epilepsies (GGE, red) datasets the burden analysis without genomic rearrangement hotspots loci consideration[14] is shown using the following microdeletion sets: All microdeletions; Overlapping at least one: CNV constrained gene, Neuro-Developmental gene[28], ASD-Related gene[29], developmental disorders[30] genes and loss-of-function intolerant genes[31]. The effect size observed (OR), confidence interval (C.I., horizontal lines) and multiple testing corrected p value obtained is shown for each dataset. Triangles denote if the signal is nominally significant. C.I. above 9 are shown in numbers.

### Microdeletions gene content: Epilepsy candidate gene identification outside hotspot loci

To extract additional epilepsy genes of interest overlapped by microdeletions outside hotspot loci (already examined in[18]) we generated a short list of potential epilepsy candidate genes that can be followed up in future studies. First, we performed a gene-oriented burden analysis outside hotspot loci comparing the number of cases carrying a microdeletion within a gene against the corresponding number of controls. We detected nominal association for *NRXN1* and *RBFOX1* genes (Table 2, p nominal = 0.019, No. of cases = 3, No. controls = 0). In this regard, due to the low frequency of microdeletion events, extraordinary large sample sizes are required to identify significant enrichment for a particular gene. Secondly, we report genes outside hotspot loci boundaries, affected more than once by microdeletions in cases and not in controls (e.g case-only genes). Candidate genes fulfilling these filtering criteria are resumed in Table 2. Finally, we also highlight genes with at least one case affected by a microdeletion belonging to the DDG2P gene list for developmental disorders[30], namely *SKI*, *KCNA2*, *GCH1,* and *DVL1.*

**Table 2.** Candidate gene identification outside hotspot loci.

| Nominally Associated Gene | Chr | Start | End | Patients (N=2,454) | Ctrl (N=6,746) | P-Value | Constrained Gene+ |
| --- | --- | --- | --- | --- | --- | --- | --- |
| RBFOX1 | 16 | 6,069,131 | 7,763,340 | 3 (AFE, AFE, GGE) | 0 | 0.0190 | No |
| NRXN1 | 2 | 50,145,642 | 51,259,674 | 3 (GGE, GGE, RE) | 0 | 0.0190 | No |
| **Candidate Gene(s)** |  |  |  |  |  |  |  |
| LOC102723362 | 2 | 22,759,350 | 22,761,159 | 2 (GGE, RE) | 0 | 0.0711 | Yes |
| PCDH7 | 4 | 30,722,036 | 31,148,423 | 2 (AFE, GGE) | 0 | 0.0711 | No |
| PACRG | 6 | 163,148,163 | 163,736,524 | 2 (GGE, AFE) | 0 | 0.0711 | Yes |
| ADGRB1,<br>TSNARE1,<br>LINC00051,<br>MIR1302-7,<br>MIR4472-1,<br>MIR4539 | 8 | 142,86,602 | 143,626,368 | 2 (GGE, GGE) | 0 | 0.0711 | No,<br>No,<br>Yes,<br>Yes,<br>Yes,<br>No |
| LOC101928137 | 12 | 73,552,969 | 73,602,097 | 2 (GGE, GGE) | 0 | 0.0711 | Yes |

Single gene and candidate gene enrichment analysis. Associated genes: Genes identified due to nominal enrichment in patients. Candidate genes: Epilepsy candidate genes deleted at least twice in patients and not deleted in the control cohort. +Constrained gene according to the CNV map of Healthy individuals[8].

Overall, *NRXN1*, *PCDH7, TSNARE1* and *RBFOX1*, have been reported previously associated with GGE[18 22]. While microdeletions overlapping *RBFOX1* were expected to be present also in the AFE cohort based on the original study [20], we also identified additional microdeletions overlapping *NRXN1* and *PCDH7* carried by RE and AFE patients too. Interestingly, CNVs overlapping *LOC102723362* have been reported in patients with neurodevelopmental phenotypes, including autism and intellectual disability (DECIPHER entries 278594, 260750 and 249773, database version 9.3)[30]. Microdeletions overlapping *PACRG* were observed in patients with AFE and GGE. Furthermore, two patients with GGE had microdeletions overlapping the *ADGRB1* as well as three non-coding genes (*LINC00051*, *MIR1302-7*, *MIR4472-1* and *MIR4539*). For *LOC102723362*, *PACRG,* and *LOC101928137,* no microdeletions were reported in the curated CNV map of healthy individuals[8] in the Database of Genomic Variants (Table 2).

### Microdeletions gene content: Tissue specific expression and network analysis of candidate genes

To further evaluate the plausible involvement of the selected candidate genes in neuronal processes and epilepsy, we performed a global gene set enrichment analysis using expression data from the GTEx portal (http://www.gtexportal.org/home/; version3). We compared the expression patterns of the 12 genes deleted more than once in patients (Table 2) versus the 92 genes deleted more than once in controls. While the analysis shows an emerging pattern for brain tissues not observed in controls, after correction for multiple testing, the observation was not significant (figure 2). Finally, to evaluate the entire pool of genes only deleted in patients in a network context we used the Disease Association Protein-Protein Link Evaluator (DAPPLE)[33] to assess if these genes form protein-protein interactions networks that have more interconnections than expected. We ran the analysis for each epilepsy syndrome separately, with only the GGE syndrome being significantly interconnected (figure 3; P = 0.025). The only cluster with multiple significant genes consists of *GRM1*, *ARC*, *PLCB1*, *MAPK3* and *PLA2G10*. Interestingly, four of these genes (*GRM1*, *PLCB1*, *MAPK3* and *PLA2G10*) are members of the KEGG Long Term Depression gene set (hsa04730, nominal P = 8.45 × 10^-7^), and three (*GRM1*, *PLCB1*, *MAPK3*) in the KEGG Long Term potentiation network (hsa04720, nominal P = 2.06 × 10^-4^).

**Figure 2.**
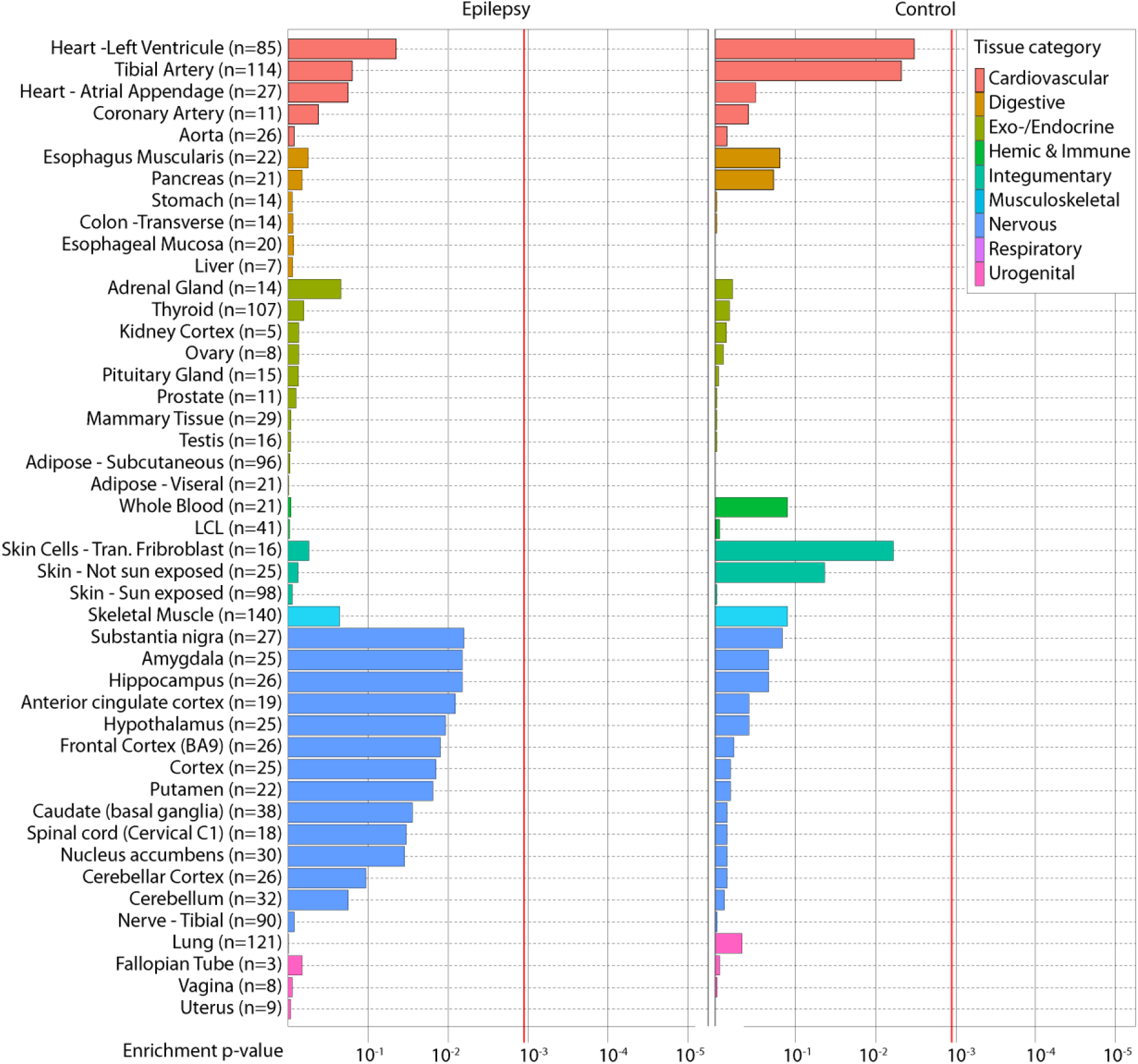
Tissue specific gene expression of epilepsy patient deleted genes. Tissue enrichment analysis using publicly available expression data from pilot phase of the Genotype-Tissue Expression project (GTEx), version 3 (see Methods). Overall, 45 individual tissues were assayed and grouped in to nine categories by color (upper-right box). Left panel: Results for genes identified to be deleted more than once in patients and not in controls (n = 12). Right panel: Results for genes deleted more than once in controls and not in patients (n = 96). For both analysis the significance threshold is denoted by vertical red line (P=1.90 × 10^-3^).

**Figure 3.**
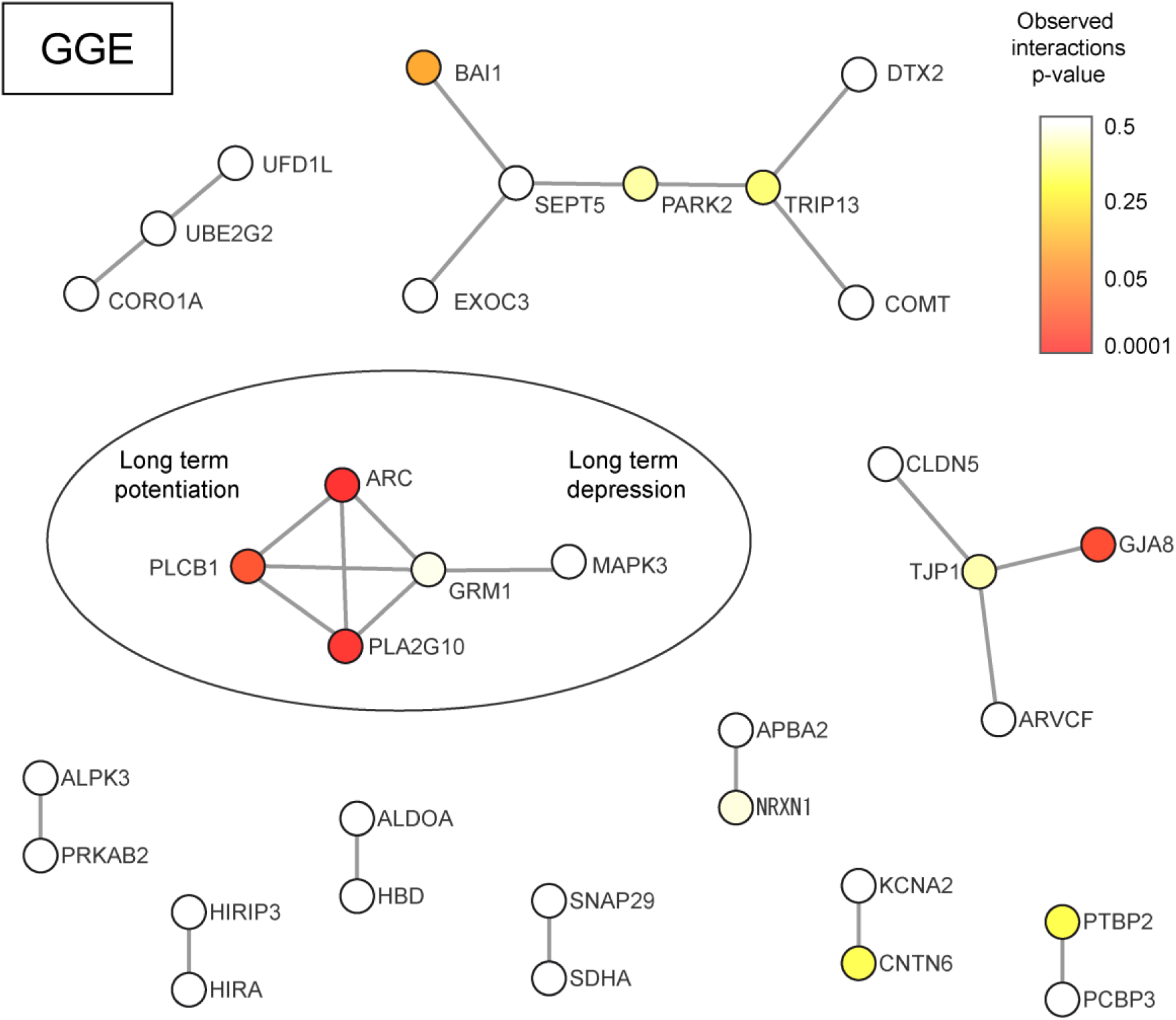
Protein-protein interaction network of GGE exclusively deleted genes. Total interacting structures found in GGE exclusively deleted genes (not deleted in controls) are shown. Each gene (node) in the network is colored based on the significance of having more-than-expected edges (interactions) following the p-value legend provided at the upper-right corner. Significant cluster is enclosed within a circle.

## DISCUSION

We have previously shown in GGE patients an enriched burden of microdeletions overlapping neurodevelopmental genes in comparison to controls and that the signal was particularly concentrated on seven hotspot loci[18]. In the present work we moved one step further and examined if this burden extends broadly (combined analysis) and/or specifically to other types of epilepsies (sub-type analyses). While we were able to replicate the original GGE signal, novel analyses were introduced to further characterize the GGE burden, as well as in the RE and AFE cohorts, including: the evaluation of twelve hotspot loci, microdeletions overlapping constrained genes, neurodevelopmental disorders genes (DDG2P) and Loss of Function Intolerant (LoF) genes. Moreover, when focusing on genes exclusively deleted in cases we were able to extract additional genes with plausible pathogenic behavior.

### Burden of microdeletions in epilepsy patients

In the combined analysis, we found a 1.39 fold excess of patients with microdeletions compared to controls, which translates in to 4.85% of epilepsy patients carrying at least one microdeletion compared to 3.47% of controls. At this level, microdeletions exhibit a mild impact on the overall genetic susceptibility towards common epilepsy types, which is particularly concentrated on GGE. The spectrum of 134 microdeletions identified in patients of the combined epilepsy analysis contained a high proportion of recurrent microdeletions at genomic rearrangement hotspot loci (32.34 %), again with the highest fraction coming from GGE patients (90.47 %). Although, we controlled for array differences (Affymetrix vs Illumina) in the combined analysis, we can not be certain that platform bias was completely removed and thus conclude that RE or AFE do not have additional microdeletion enrichment to contribute, especially considering that individuals with microdeletions only account for a minor proportion of patients. Similarly, in the sub-type specific analysis we only compared platform matched cases with controls, nevertheless the complete contribution of microdeletions to the presentation of each epilepsy sub-type could be underestimated since our analysis was restricted only to high confidence calls and large microdeletions based on high-throughput genotyping. Furthermore, our cohort size, in particular the RE cohort, is still too small to identify microdeletions with small disease risk. We also acknowledge the high heterogeneity of the AFE cohort (see definition in Supplementary Information) where the general pool of patients with focal epilepsy could overshadow specific and rare pathogenic events. Larger samples size, prospective studies and deep phenotyping will allow the evaluation of other rare variants in the presence of microdeletions that could explain the specific outcomes commonly observed in neurodevelopmental disorders. Still, considering the aforementioned caveats, our results enable of us to elucidate the followings topics.

### The landscape of microdeletions overlapping and not overlapping hotspot loci is epilepsy subtype-specific

In line with the original GGE study [18], the strongest combined enrichment was found for microdeletions overlapping known microdeletion syndrome hotspot loci. The level of association did not increase with the inclusion of additional 5 hotspot loci to the analysis. These microdeletions are commonly found in patients with other highly comorbid traits such as ASD, ID and SCZ giving rise to a complex network of neurodevelopmental phenotypes [14 16]. The specific effect of each of the hotspot loci evaluated has been difficult to determine and certainly the outcome of microdeletions overlapping these regions is not epilepsy exclusive.

When these regions are considered, sub-type analyses show that for the GGE cohort, the strongest signal falls within genes overlapping neurodevelopmental genes (as previously shown in [18]). In contrast, for RE the frequency of microdeletions was similar to controls in all subsets exanimated, with the exception of a modest enrichment and a large OR observed for microdeletions overlapping ASD-related genes (Supplementary figure S2). While the latter observations was not significant we can not rule out that future larger studies will identify a microdeletion burden. The AFE sub-type analysis did not show enrichment for any of the investigated microdeletion subsets with hotspots included. In this regard, we observed that the contribution of these regions is compelling and epilepsy subtype-specific. Notably, by removing them from the analysis, the only significant enrichment that remained was for genes associated with neuronal development in the GGE sub-type (Figure 1).

The enrichment for GGE with hotspot loci has previously been shown to be more significant in patients with GGE and ID[34]. In this regard, in contrast to severe neurodevelopmental disorders with or without seizures[35] the microdeletion burden for common epilepsy patients with normal social and intellectual skills (i.e. high-functioning) is expected to be modest, thus we cannot rule out that GGE patients in our cohort with hotspot microdeletions represent those at the lower boundaries of the IQ distribution in the general population. It has previously been shown that CNVs at hotspot loci affect cognition in patients and controls[36 37]. Given that the majority of the identified hotspot CNVs have been also associated with intellectual disability, well phenotyped cohorts are needed in future studies to investigate whether seizures are also an independent consequence of these particular microdeletions.

### Established disease and candidate genes only deleted in patients outside of microdeletion hotspots

Four patients carried microdeletions overlapping the established developmental disorder genes *SKI*, *KCNA2*, *GCH1* and *DVL1.* As an additional support for haploinsufficiency of these genes, all show in >60,000 population controls depletion for loss-of-function variants (http://exac.broadinstitute.org). We identified nominal enrichment for the gene encoding the adhesion molecule neurexin 1 (*NRXN1*) and the splicing regulator RNA-binding protein fox-1 homolog *(RBFOX1)*, which were previously implicated in epilepsy and neurodevelopmental disorders[20 22 23 38 39]. To identify plausible candidate genes for epilepsy with potentially large effects, we selected genes exclusively overlapped by at least two microdeletions in epilepsy patients. We identified 10 candidate genes at seven loci. These autosomal microdeletions involved several genes previously implicated in epilepsy and neurodevelopmental disorders. Specifically, the genes encoding T-SNARE Domain Containing 1 (*TSNARE1*) and protocadherin 7 (*PCDH7*) have been highlighted previously in our GGE microdeletion analysis[18] (1,366 patients and 5,234 controls) which was entirely integrated in the present study. Furthermore, with the analysis of other types of epilepsies we detected one RE patient with a partial *NRXN1* microdeletion and one AFE patient with a complete *PCDH7* deletion. These observations suggest their role as broader epilepsy risk factors rather than syndrome specific variants of high effect. Furthermore, our gene-centric (compared to microdeletion-centric) analysis could narrow down four large microdeletions to *PCDH7*, *PACRG*, *LOC102723362* and *LOC101928137* as the only remaining genes not deleted in controls respectively. We acknowledge the limitations of this analysis since we do not have the power to assign a meaningful p-value to these detected genes.

### Expression and network analysis

In the expression analysis of candidate genes we did not observe significant brain tissue enrichment, probably because the number of included genes was small and not all of them may be involved in epilepsy. While global or individual-gene brain expression patterns would have been informative, the results are not conclusive and thus we cannot rule out candidates based on gene expression filtering. The network analysis resulted in significant interconnection only for the GGE syndrome. The likely reason for this is the difference in the number of regions between the syndromes rather than a difference in the underlying biology. These results are encouraging, considering that we use non cell-type specific networks and did not filter the network based on tissue-specific gene expression. The enriched KEGG Long Term Depression (hsa04730) and Long Term Potentiation (hsa04720) networks represent plausible neuronal enriched networks in epilepsy patients.

In summary, we show that the microdeletion enrichment in epilepsy patients is focused towards genes involved in neurodevelopmental processes. Patients with GGE syndrome exhibit the highest microdeletion frequency, especially at hotspot loci. Apart from these loci ultra-rare heterogeneous deletions contribute significantly to GGE whereas microdeletion frequency and distribution in AFE is indistinguishable from controls. The RE cohort is the smallest and therefore has the lowest statistical power for association discovery. However, the RE cohort shows nominal enrichment for hotspot loci microdeletions. There was some support for ultra-rare microdeletions as plausible epilepsy candidate genes. Our study demonstrates, that the contribution of microdeletions in common epilepsies is subtype specific. With increasing cohort sizes, the genetic architecture of the epilepsies and the contribution of microdeletions will become more evident.

Despite these differences, candidate genes can be found commonly deleted in more than one epilepsy type. Thus, the present findings contribute to our understanding of the structural genetic architecture of epilepsies from an overall and sub-type specific perspective.

## ACKNOWLEGDMENTS

We thank all the participants and their families. We thank the EuroEPINOMICS-RES consortium and the The Italian League against Epilepsy (LICE). A full list of contributors and associated centers is provided in the Supplementary Information File. This study makes use of data generated by the DECIPHER community. A full list of canters who contributed to the generation of the data is available from http://decipher.sanger.ac.uk and via email from. Funding for the project was provided by the Wellcome Trust.

## CONTRIBUTOR

Design and coordination of the study: E. P-P, I.H. and D. L. Performed microdeletions statistical analysis: E. P-P, P. G., A. G., K. P., E. S. and D. P. H. Performed GTex expression analysis: V. A, A. B. Performed Network analysis: H. H.; Contributing genetic and/or phenotypic data: M. R. T., E. M. R., P. N., H. L., F. Z, A. J. B., F. R., E. P., F. Z. and Y. G. W.; Writing of the manuscript: E. P-P, I.H., and D. L. Revision of the manuscript with important intellectual content: J.F.M., G.D.F, K. M. K., B. A. N., P. N. and D. L.

